# cAMP signaling mediates behavioral flexibility and consolidation of social status in *Drosophila* aggression

**DOI:** 10.1101/089979

**Authors:** Nitin Singh Chouhan, Krithika Mohan, Aurnab Ghose

## Abstract

Social rituals, like male-male aggression in *Drosophila*, are often stereotyped and the component behavioral patterns modular. The likelihood of transition from one behavioral pattern to another is malleable by experience and confers flexibility to the behavioral repertoire. Experience-dependent modification of innate aggressive behavior in flies alters fighting strategies during fights and establishes dominant-subordinate relationships. Dominance hierarchies resulting from agonistic encounters are consolidated to longer lasting social status-dependent behavioral modifications resulting in a robust loser effect.

We show that cyclic adenosine monophosphate (cAMP) dynamics regulated by the calcium/calmodulin-dependent adenylyl cyclase, Rut and the cAMP phosphodiesterase, Dnc but not the Amn gene product, in specific neuronal groups of the mushroom body and central complex, mediate behavioral plasticity necessary to establish dominant- subordinate relationships. *rut* and *dnc* mutant flies are unable to alter fighting strategies and establish dominance relationships during agonistic interactions. This real-time flexibility during a fight is independent of changes in aggression levels. Longer-term consolidation of social status in the form of a loser effect, however, requires additional *Amn*-dependent inputs to cAMP signaling and involves a circuit-level association between the α/β and γ neurons of the mushroom body.

Our findings implicate cAMP signaling in mediating plasticity of behavioral patterns in aggressive behavior and in the generation of a temporally stable memory trace that manifests as a loser effect.

**SUMMARY STATEMENT:** Phasic recruitment of different cAMP signaling modalities in specific neuronal groups lead to the formation of temporally distinct components of learning and memory in fly aggression.

## INTRODUCTION

Aggression is a social behavior involving competitive interactions over resources to ensure reproductive success and survival. Conspecific agonistic interactions may result in dominance hierarchies, where those at higher levels have better access to resources. The social ritual of *Drosophila* aggression is modular and comprises of stereotyped behavioral patterns analogous to a sequence of fixed action patterns. These sequences are present in full complexity in socially naive animals and appear to be pre-wired in the nervous system (Chen et al., 2002; Hoopfer, 2016; Lim et al., 2014). However, during aggressive encounters, experience can alter the likelihood of transitions between specific patterns and lead to experience-dependent plasticity (Trannoy et al., 2016; Yurkovic et al., 2006). Initial agonistic interaction between naïve flies constitutes a conditioning phase where male flies employ a combination of offensive and defensive fighting strategies and display real time, experience-dependent plasticity to establish dominant or subordinate status (Chen et al., 2002; Yurkovic et al., 2006). In flies, the learned subordinate social status is consolidated into a long lasting loser effect, resulting in an increased probability of losing agonistic encounters against familiar as well as unfamiliar opponents (Trannoy and Kravitz, 2016; Trannoy et al., 2016; Yurkovic et al., 2006).

Aggression is a complex social behavior influenced by a combinatorial interplay between genetic factors, environmental cues and experience. Population level selection for elevated aggression has implicated several genes, which show significant changes in their expression (Dierick and Greenspan, 2006; Edwards et al., 2009; Wang et al., 2008). However, a single social defeat can lead to the development of a robust loser effect in such hyperaggressive lines, underscoring the modulatory role of experience in relation to intrinsic abilities (Penn et al., 2010). While regulatory activities of pheromones, neuromodulatory agents and neurotransmitters associated with aggression levels are well documented (Alekseyenko et al., 2014; Alekseyenko et al., 2013; Andrews et al., 2014; Certel et al., 2010; Dierick and Greenspan, 2007; Hoopfer, 2016; Hoyer et al., 2008; Kohl et al., 2015; Luo et al., 2014; Wang and Anderson, 2010; Wang et al., 2011; Yuan et al., 2014), little is known about the neurogenetic underpinnings of the relevant learning and memory components.

Social experience significantly alters the activity of flies and involves both classical and operant conditioning components (Kamyshev et al., 2002). Experience-dependent behavioral plasticity has also been observed in courtship conditioning assays where males, after an unsuccessful mating experience, modulate their courtship behavior (Siegel and Hall, 1979). Learning and memory in a social environment are likely to be influenced by several factors, including the combinatorial inputs from multiple sensory modalities. Consequently, central integration and interpretation of complex input modalities are likely to involve coordination between multiple neurogenetic circuits.

Previous studies have established the centrality of the cyclic adenosine monophosphate (cAMP) pathway in the formation of operant and classical conditioned memories and in the integration of sensory inputs (Bragina and Kamyshev, 2003; Brembs, 2003; Busto et al., 2010; Gailey et al., 1984). Several mutations associated with defective learning and memory have been mapped to the genetic loci of cAMP pathway components (Dudai et al., 1976; Feany and Quinn, 1995; Levin et al., 1992; Livingstone et al., 1984). In this study, we implicate the cAMP second messenger pathway in behavioral plasticity during aggressive encounters and in the development of the loser effect. *rutabaga* (calcium- calmodulin-dependent adenylyl cyclase; Rut) and *dunce* (cAMP phosphodiesterase; Dnc) mutants show compromised behavioral flexibility during agonistic encounters and are unable to establish dominance hierarchies. *amnesiac* (predicted to code for three putative neuropeptides; Amn) mutants, although competent in modifying behavioral patterns during fights and establishing hierarchies, show no loser effect. These studies demonstrate that *Rut* in specific neural circuits mediates behavioral flexibility leading to the establishment of dominance relationships and in the long-term consolidation of social status.

## MATERIALS AND METHODS

### Fly stocks and maintenance

Canton S (CS), *rut^2080^* and *amn^c651^* lines were obtained from Dr R. Strauss, University of Mainz, Germany. *dnc^1^, rut^2080^;UAS-Rut* and all Gal4 driver lines were obtained from the stock center at Bloomington, USA. All fly lines were backcrossed for at least nine generations. The autosomes of the X-linked cAMP pathway mutant alleles used were equilibrated to that of the control strain. Stocks were maintained at 25°C, 60% humidity and a 14h:10h∷light:dark cycle on standard food.

### Analysis of aggressive behavior

Freshly eclosed flies were isolated and kept in social isolation for a period of 4-5 days before testing. Acrylic paint marks on the upper thoracic region were used to identify individuals. Male-male aggression assays were conducted as described earlier (Chen et al., 2002) with the modification that they were conducted in a six-well chamber. All experiments were conducted between Zeitgeber Time (ZT) 0 and ZT 6 unless otherwise specified. A food cup with yeast paste and a headless female was placed inside a six- well plate chamber. A pair of marked, un-anesthetized, age- and size- matched male flies was introduced into the chamber through gentle aspiration. All fights were conducted at 25°C and 60% humidity and recorded using a Sony DCR-SR47E/S video camera at 50 Hz. Fights between socially naïve flies involved three phases. In the ‘fight phase’ a pair of naïve flies was allowed to fight for 60 min, this was followed by a ‘rest phase’ of 60 min where flies were returned to their original food vials and finally a ‘test phase’ where previously matched flies fought against unfamiliar, naïve opponents for 60 min. SONY PMB software on Windows OS was used for video playback and the fights were manually curated. In all cases the analyzer was blind to the genotype of the fly. Fights were analyzed on basis of encounters that involved physical interactions and lasted at least 3 seconds. Encounters were considered separate if the time interval between them exceeded 2 seconds. Winners and losers were designated based on an ethologically characterized three lunge-three retreat rule (developed in (Yurkovic et al., 2006)). Lunge is defined as a maneuver where a fly rears up on its hind legs and collapses on the opponent (Chen et al., 2002; Zwarts et al., 2012). A retreat occurs when a fly, in response to an offensive action, runs/flies away from the opponent or the food cup (territory) (Chen et al., 2002; Zwarts et al., 2012). Within a trial, a fly was assigned a status of ‘winner’ if it used three continuous lunges (without any interim retreats) and its opponent executed three corresponding retreats (with no intervening lunges) (Yurkovic et al., 2006). The latter was designated as a ‘loser’.

We analyzed various parameters to investigate aggression in flies across genotypes. They include: 1. Encounter frequency (encounters per minute measured as the total number of encounters divided by total time of fighting); 2. Aggression vigor index (the fraction of time spent fighting in the first 10 min following the commencement of the first encounter). 3. Latency to engage in an encounter (time in seconds from the start of the fight to initiate the first agonistic interaction that lasts for at least 3 seconds). In the second fights, the ‘Draw’ outcomes, where 3 Lunge- 3 Retreat rule was not satisfied, were further divided into three categories based on Penn *et al*. (2010). This categorization was based on usage of lunges and retreats by experienced loser flies against a naive opponent. ‘High intensity’ draws consisted of experienced loser flies predominantly using lunges while ‘low intensity’ draws included usage of retreats. Fights in which the flies did not engage in agonistic interactions were considered as ‘no intensity’ and categorized as draws. An experienced loser fly may not readily engage in agonistic interactions against naïve opponents. This may preclude escalation during fights and result in higher number of draws. Thus ‘high/low intensity’ categorization facilitates better assessment of the status-dependent behavioral changes manifested as the loser effect.

We devised the loser index in order to assess the ability of flies to demonstrate the experience dependent loser effect. The loser index was calculated as a difference between the numbers of encounters lost to the encounters won divided by the total number of encounters in the second fight. Within an encounter, if a fly uses aggressive actions like lunging, tussling (tugging and pulling the body of the opponent), boxing (rearing up on hind legs and striking the opponent with forelegs), holding (rearing up on hind legs and holding the other fly’s abdomen), chasing (running after the opponent) or fencing (extending its leg forward and pushing the other fly) (Chen et al., 2002; Zwarts et al., 2012) and the other fly responds with a retreat then the former fly is a ‘winner’ and the latter a ‘loser’ in that encounter.

### Analysis of locomotor behavior

Locomotor behavior was analyzed using a negative geotaxis assay as described before (Ali et al., 2011). A group of ten flies, age and size matched, were introduced into a food vial one day prior to testing. The flies were placed in two head to head joined empty food vials with a distance of 8 cm marked on the lower vial. Following a gentle tap to get all the flies to the base of the vial, the number of flies able to climb above the 8 cm mark in 10 seconds was scored. The experiment was repeated 10 times for each group and the average pass rate calculated. Three experimental replicates were carried out for each genotype.

### Statistical analysis

Videotapes were analyzed and each encounter was scored for all fighting strategies and documented on spreadsheets. All statistical analysis was performed using Statistica 8.0 and GraphPad Prism 6.0 statistical software. Various statistical tests were employed to facilitate our analysis including Chi-square test, one-factor ANOVA and two-factor repeated measures ANOVA followed by posthoc Tukey’s or Dunnett’s multiple comparisons tests. Specific information on statistics used in each experiment is included in the figure legends.

## RESULTS

### The cAMP pathway mediates establishment of hierarchies within a fight

Pairs of male flies demonstrate stereotyped aggressive behavior towards each other while competing for resources like food or females. These dyadic interactions begin with low aggression maneuvers like wing threat and fencing, which then escalate to high aggression strategies like lunging and tussling. This escalation is a consequence of experience-dependent behavioral transitions, with one fly demonstrating offensive strategies with increasing frequency against an opponent fly employing more and more defensive strategies. These real time behavioral modifications on shorter time scales are stable within a fight and facilitate formation of social dominance hierarchies. We have assigned dominance relationships using a previously developed three lunge-three retreat rule (see Materials and Methods and (Yurkovic et al., 2006)). In wild-type Canton S (CS) flies, 85% of the fights produced dominance relationships (Figure 1A). Prior investigations have implicated the cAMP pathway in the development of learning and memory in multiple conditioning paradigms. cAMP synthesis is regulated by the enzymatic activity of the *Rut* encoded adenylyl cyclase that catalyzes the conversion of ATP to cAMP (Levin et al., 1992; Livingstone et al., 1984). *Rut* has been implicated as a biochemical coincidence detector necessary for the formation of short term memory (STM) in olfactory conditioning assays (Dudai et al., 1988; McGuire et al., 2005). In contrast to CS flies, *rut^2080^* mutant flies were unable to form dominance relationships with all their fights ending in draws (Figure 1A; *P* < 0.001). cAMP phosphodiesterase activity encoded by *Dnc* negatively regulates cAMP levels (Davis and Kiger, 1981). Consistent with the deregulation of cAMP dynamics, *dnc^1^* flies display attenuated formation of dominance relationships with only 67% of the fights resulting in wins/losses (Figure 1A; *P* < 0.01). The *Amn* gene product is thought to encode for neuropeptide/s and is known to modulate cAMP signaling (Feany and Quinn, 1995; Moore et al., 1998) However, *amn^C651^* mutant flies displayed wild-type levels of dominance hierarchies (Figure 1A; 82% of the fights, *P* > 0.05).

**Figure 1.**
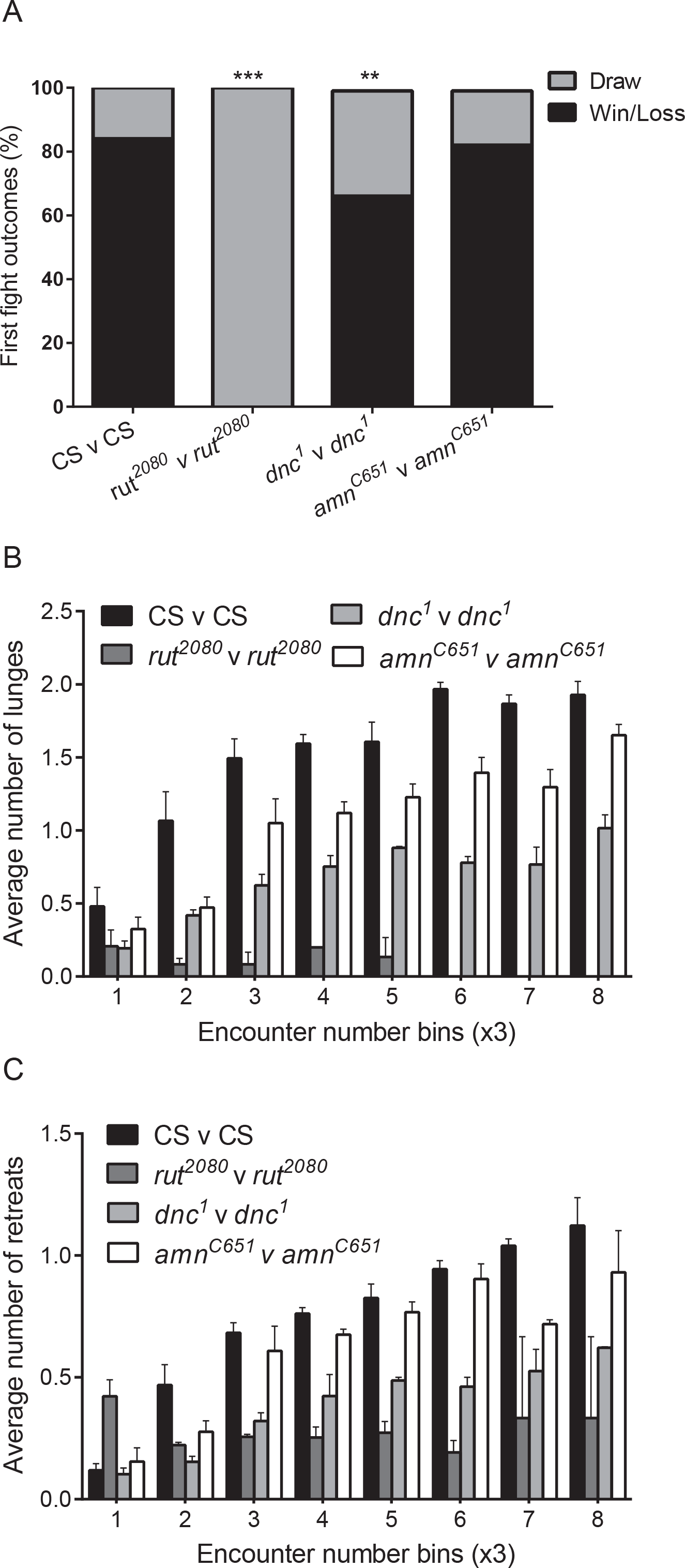
cAMP signaling is required for establishing dominance relationships. (*A*) CS and *amn^C651^* flies form stable dominance relationships (win/loss) in 85% and 82% in their first fights, respectively. *rut^2080^* displayed no dominance structures while *dnc^1^* showed dominance in 67% of their fights. (Two-tailed Chi-square test; ** *P* < 0.01, *** *P* < 0.001, n=30). *, indicates comparison with CS. (*B*) CS winner flies progressively increase lunging. A similar trend was seen with amn^C651^flies (*P* > 0.05). In contrast, *rut^2080^* (*P* < 0.001) and *dnc^1^* (*P* < 0.01) flies demonstrated significantly deficient lunging trends. (Two factor repeated measures ANOVA followed by post hoc Tukey’s multiple comparison test, ‘*P*’ is the interaction term, mean ± SEM, n=30). (*C*) Both CS and *amn^C651^*loser flies (*P* > 0.05) showed progressive increase in retreats while this was significantly compromised in *rut^2080^ (P* < 0.001) and *dnc^1^* (*P* < 0.01). *dnc^1^* mutants, although compromised in modifying fighting strategies compared to CS flies, demonstrate better trends compared to *rut^2080^* mutants (*P* < 0.001 for lunges; *P* < 0.01 for retreats) (Two factor repeated measures ANOVA followed by post hoc Tukey’s multiple comparison test, ‘*P*’ is the interaction term, mean ± SEM, n=30). Total number of lunges/retreats in every three successive encounters is analyzed and their mean is reported. SEM: Standard error of the mean.

For further analysis of experience dependent behavioral modifications during a fight, we focused on lunges and retreats. These strategies are commonly used offensive (lunges) and defensive (retreats) maneuvers and previous work have indicated status dependent modifications in their usage by winner/loser flies (Yurkovic et al., 2006). Behavioral modulation was assessed by scoring numbers of lunges/retreats in first 25 encounters in a fight. In line with previous observations, agonistic interactions resulted in an escalation of aggression (Trannoy et al., 2016; Yurkovic et al., 2006). In aggressive encounters between CS flies, the ultimate winners (as established by the 3 lunge-3 retreat rule; see Materials and Methods) progressively increased the deployment of lunges while the losers increasingly adapted to the retreating behavior (Figure 1B, C). In contrast, *rut^2080^* mutant flies were unable to demonstrate any behavioral transitions pertaining to lunges and retreats (Figure 1B, C; *P* < 0.001). *dnc^1^* flies were not only inefficient in establishing hierarchies but also displayed compromised behavioral plasticity with significantly reduced ability to modify the frequency of lunges and retreats compared to CS animals (Figure 1B, C; *P* < 0.001). *amn^C651^* flies, which are competent in establishing hierarchies, were able to modify the usage of offensive and defensive strategies as well as CS flies (Figure 1B, C; *P* > 0.05). These results suggest a correlation between experience dependent behavioral plasticity and formation of dominance hierarchies following aggressive encounters.

No significant differences in locomotor activity between wild type and mutant flies were found using a negative geotaxis assay (see Materials and Methods), thus ruling out the possibility of overt motor deficits in these lines (Figure 2A; *P* > 0.05). Multiple parameters were evaluated to assess the aggression levels of the wild type and mutant lines. Encounter frequency was significantly lower in *rut^2080^* mutants compared to CS flies but *dnc^1^* and *amn^C651^* mutants were comparable to wild-type flies (Figure 2B; *P_ru_t* < 0.05, *P_dnc_* and *P_amn_* > 0.05). Aggression vigor index was compromised in all the mutant lines compared to CS but was not significantly different between *rut^2080^, dnc^1^* and *amn^C651^* flies (Figure 2C; *P_rut_* < 0.001, *P_dnc_* and *P_amn_* < 0.05).

**Figure 2.**
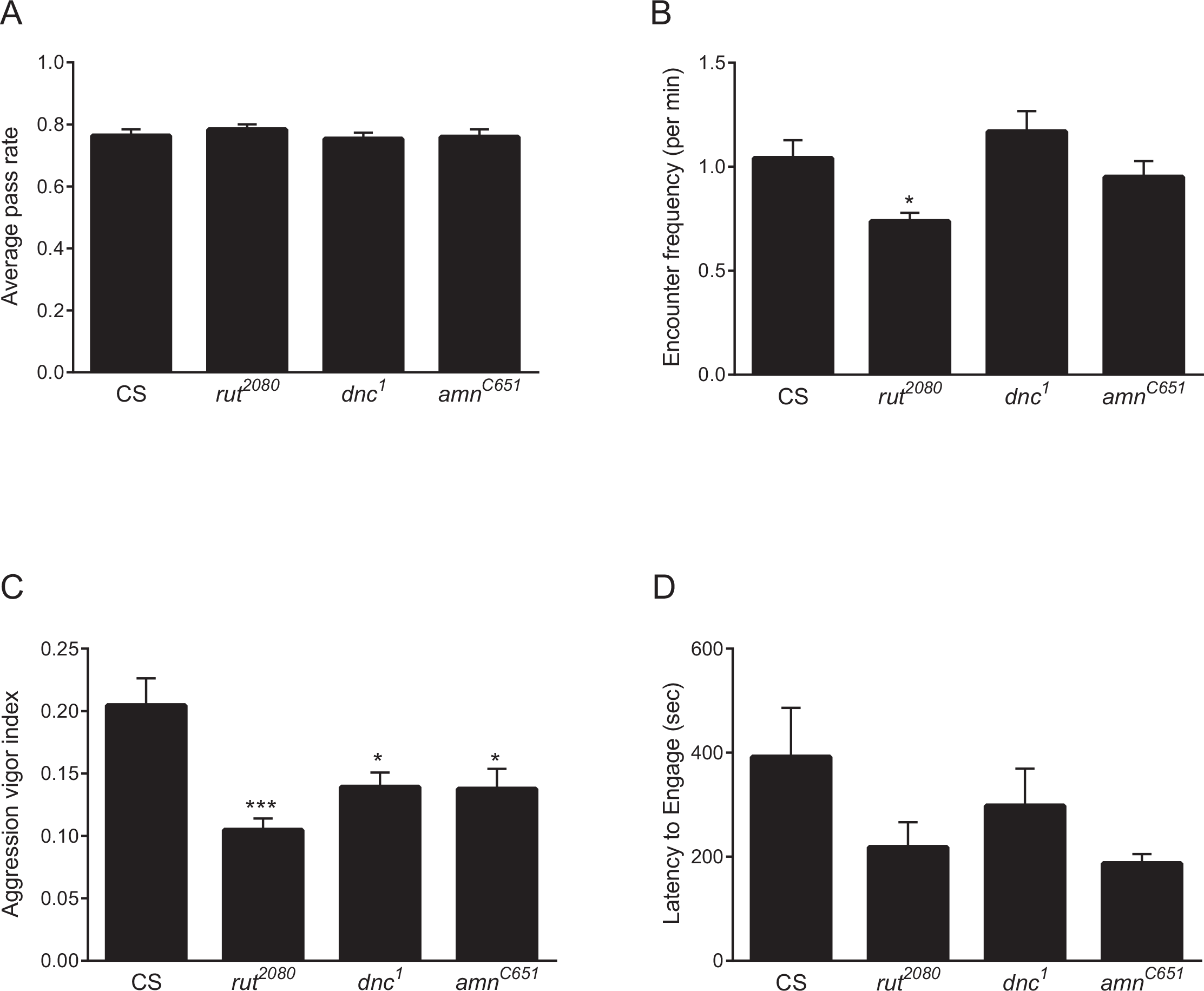
Aggression levels in cAMP pathway mutant flies. (*A*) Locomotor activity in flies was analyzed using negative geotaxis assay. No significant differences were observed between CS and cAMP pathway mutants in their average pass rate (One factor ANOVA followed by post hoc Tukey’s multiple comparison test, mean ± SEM, n=30). (*B*) Encounters per minute is lower in *rut^2080^* flies (*P* < 0.05) but is comparable in *amn^c651^* and *dnc^1^* flies compared to CS flies (*P* > 0.05) (One factor ANOVA followed by post hoc Tukey’s multiple comparison test, mean ± SEM n=30). *, indicates comparison with CS. (*C*) Aggression vigor index, defined as the fraction of time spent fighting in first 10 minutes, is significantly lower in *rut^2080^* (*P* < 0.001), *amn^c651^* (*P* < 0.05) and *dnc^1^* (*P* < 0.05) flies in comparison to CS flies (One factor ANOVA followed by post hoc Tukey’s multiple comparison test, mean ± SEM n=30). *, indicates comparison with CS. (*D*) Latency to engage in an encounter is not significantly different in *rut^2080^* (*P* > 0.05), *dnc^1^* (*P* > 0.05) and *amn^c651^* (*P* > 0.05) flies in comparison to CS flies. (One factor ANOVA followed by post hoc Tukey’s multiple comparison test, mean ± SEM, n=30). SEM: Standard error of the mean.

While our studies implicate lack of behavioral plasticity in *rut^2080^* mutants in establishing dominance hierarchies in *rut^2080^* versus *rut^2080^* fights, it is also possible that the *rut^2080^* flies are unable to execute high intensity maneuvers like lunges. In *rut-rut* fights, neither opponent can adjust their fighting patterns depending upon experience resulting in a lack of escalation beyond low intensity interactions precluding assessment of inherent behavioral changes in *rut^2080^* flies. CS flies have intact experience dependent behavioral transitions compared to *rut^2080^* mutants. Therefore, pairing CS vs *rut^2080^* may better reveal behavioral escalation during fights. CS vs *rut^2080^* fights results in significant dominance hierarchies with 37.5% of fights ending in a win or a loss, though it remains attenuated compared to CS vs CS fights (Figure 3A; *P* < 0.001). Interestingly, *rut^2080^* flies demonstrate increased lunging and retreating behavior compared to their performance in *rut-rut* fights and also win 9% of the fights against wild-type opponents (Figure 3B, C). *rut^2080^* flies, when in a *rut*-CS fight, are capable of executing and modifying the frequency of use of both lunges and retreats, although this ability remains significantly compromised compared to that in CS flies (in CS-CS fights) (Figure 3B, C; *P* < 0.001 for lunges and *P* < 0.05 for retreats). The usage of offensive and defensive strategies for *rut^2080^* flies against a wild type opponent is significantly better than *rut-rut* fights (Figure 3B, C; *P* < 0.001 for lunges and *P* < 0.01 for retreats). These results suggest that *rut^2080^* flies are capable of executing high intensity maneuvers but are compromised in modifying its frequency as well as CS flies during a fight.

**Figure 3.**
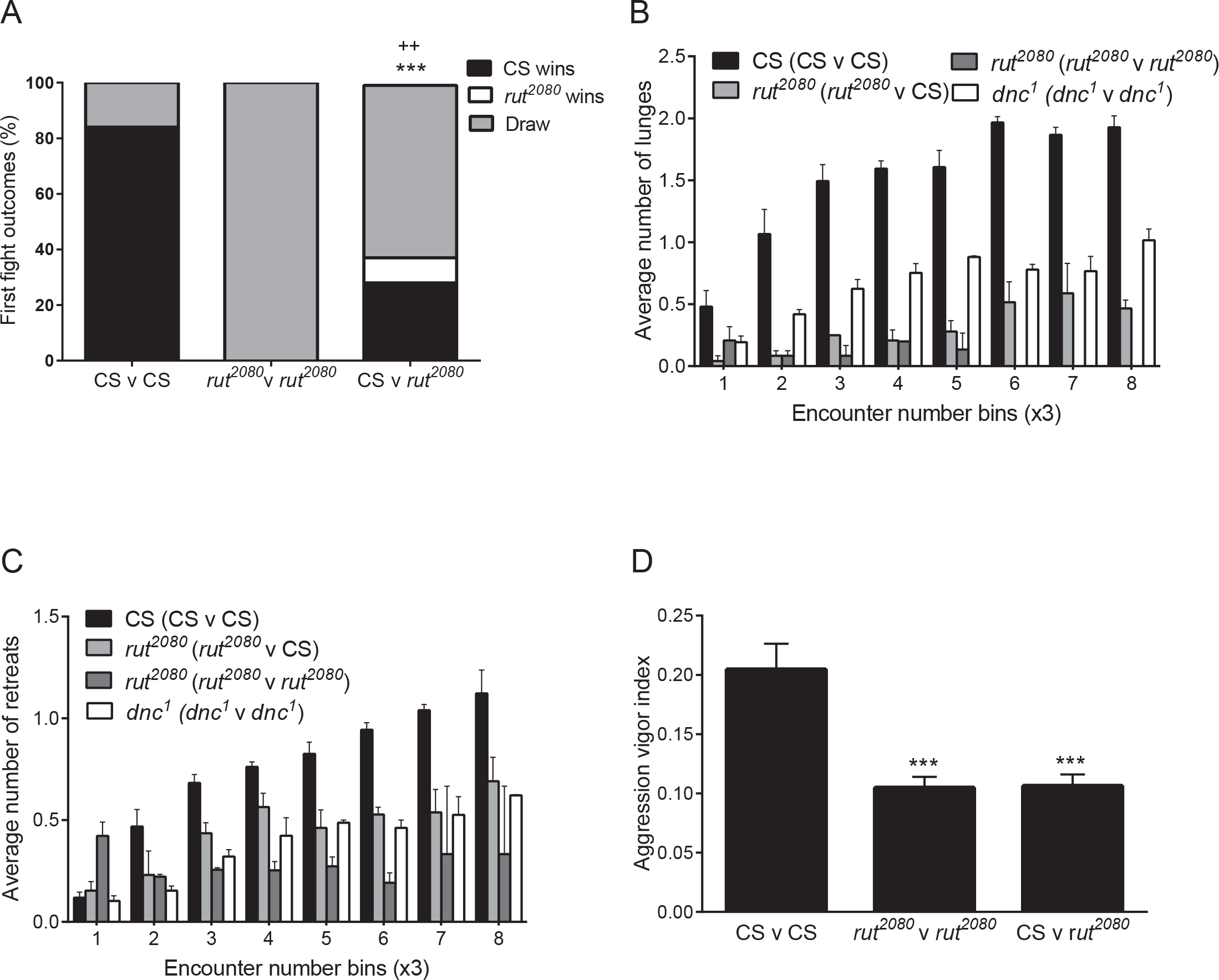
*rut^2080^* flies display improved lunging/retreating behavior against wild type opponents. (*A*) The *rut^2080^* flies show attenuated dominance structures against a wild type opponent compared to CS v CS fights (*P* < 0.001) but improved hierarchical relationships compared to *rut^2080^* v *rut^2080^* fights (*P* < 0.01) (Two-tailed Chi-square test; *** *P* < 0.001, ++ *P* < 0.01, n=25). *, indicates comparison with CS and +, with *rut^2080^*. (*B*) and (*C*) The *rut^2080^* flies against a wild type opponent demonstrate significantly better status- dependent modification of lunges/retreats compared to *rut^2080^* v *rut^2080^*fights (*P* < 0.001 for lunges and *P* < 0.01 for retreats) and is comparable to *dnc^1^* flies (*P* > 0.05). The lunge/retreat trends for *rut*^2080^flies in *rut^2080^* v CS fights are compromised compared to CS flies (*P* < 0.001 for lunges; *P* < 0.05 for retreats). (Two factor repeated measures ANOVA followed by post hoc Tukey’s multiple comparison test, ‘*P’* is the interaction term, mean ± SEM, n=20). (*D*) Aggression vigor index is significantly lower in flies engaged in *rut^2080^* v *rut^2080^* (*P* < 0.001) and CS v *rut^2080^* (*P* < 0.001) fights in comparison to flies in CS v CS fights (One factor ANOVA followed by post hoc Tukey’s multiple comparison test, mean ± SEM n=30). *, indicates comparison with flies in CS v CS fights. The data set for control CS flies, which were trained at the same time of day as experimental lines, is the same as shown in Fig 1 and 2. Total number of lunges/retreats in every three successive encounters is analyzed and their mean is reported. SEM: Standard error of the mean

Furthermore, *dnc^1^* mutants are unable to modify fighting strategies as well as CS flies but they are less compromised compared to the *rut^2080^* flies in *rut-rut* fights (Figure 1A, B, C). Consistent with the association of ability to modify fighting patterns with the establishment of dominance relationships, *dnc* mutants do establish hierarchies though less efficiently than wild-type CS flies (Figure 1A). Similarly, *rut^2080^* flies against wild type opponents demonstrate compromised plasticity and attenuated dominance structures (Figure 3A, B, C). In fact, usage of lunges/retreats by *rut^2080^* flies in *rut* – CS fights is comparable to those seen in *dnc^1^* (Figure 3B, C; *P* > 0.05 for lunges and retreats). Interestingly, the aggression vigor index is similar for *rut* - CS and *rut* - *rut* fights (Figure 3D; *P* > 0.05).

This series of experiments establish the role of the cAMP signaling in behavioral plasticity underlying modification of fighting strategies during agonistic interactions. Behavioral flexibility, rather than changes in levels of aggression, is correlated with the generation of dominance hierarchies.

### Specific neural circuits are recruited in cAMP-mediated behavioral plasticity

We next asked which components of the *Drosophila* brain were involved in the status- dependent modifications of aggressive strategies and formation of dominance relationships. As multiple sensory modalities are likely to be involved in aggressive encounters, a complex pattern of recruitment of neuronal circuits may underlie the integration of these sensory inputs (Asahina et al., 2014; Hoopfer, 2016; Hoopfer et al., 2015; Lim et al., 2014; Ramin et al., 2014; Wang and Anderson, 2010; Wang et al., 2011; Yoon et al., 2013). To test this, we used the UAS-Gal4 binary system to express the Rut gene product in restricted neuronal populations in a *rut^2080^* mutant background. We chose well-characterized Gal4 drivers that have restricted expression in defined neuronal populations of the mushroom body and central complex of the fly brain as these regions are strongly implicated as central integrators in multiple behavioral paradigms (Table 1) (Aso et al., 2009; Blum et al., 2009; Joiner and Griffith, 1999; Neuser et al., 2008; Pan et al., 2009; Torroja et al., 1999; Zars et al., 2000a; Zars et al., 2000b). All experiments were conducted at the same time of the day. The genetic rescue and the control experiments were done in parallel. Pan-neuronal expression of *Rut* using the Appl-Gal4 driver completely rescued the inability to form dominance relationships seen in *rut^2080^* mutants (Figure 4A; *P* < 0.001). Similar restoration of dominance hierarchies to wild-type levels was seen with expression limited to the a/p and y lobe neurons of the mushroom body using the c309 driver (Figure 4A; *P* < 0.001). Next, we tested a panel of Gal4 drivers expressing *Rut* in different subpopulations of neurons in the MB and the central complex of *rut^2080^* flies (Table 1). UAS-Rutabaga driven by Gal4 drivers c739, 201Y, c305a and c205, but not c819, partially rescued the inability of *rut^2080^* to form hierarchical relationships (Figure 4A; *P* < 0.001). These experiments implicate the independent involvement of the α/β, γ and α’/β’ lobes of the MB together with the F5 neurons of the fan-shaped body (FB) in neuronal processing leading to the formation of dominance. The ellipsoid body (EB) of the central complex does not appear to be involved in this function (Figure 4A; *P* > 0.05).

**Figure 4.**
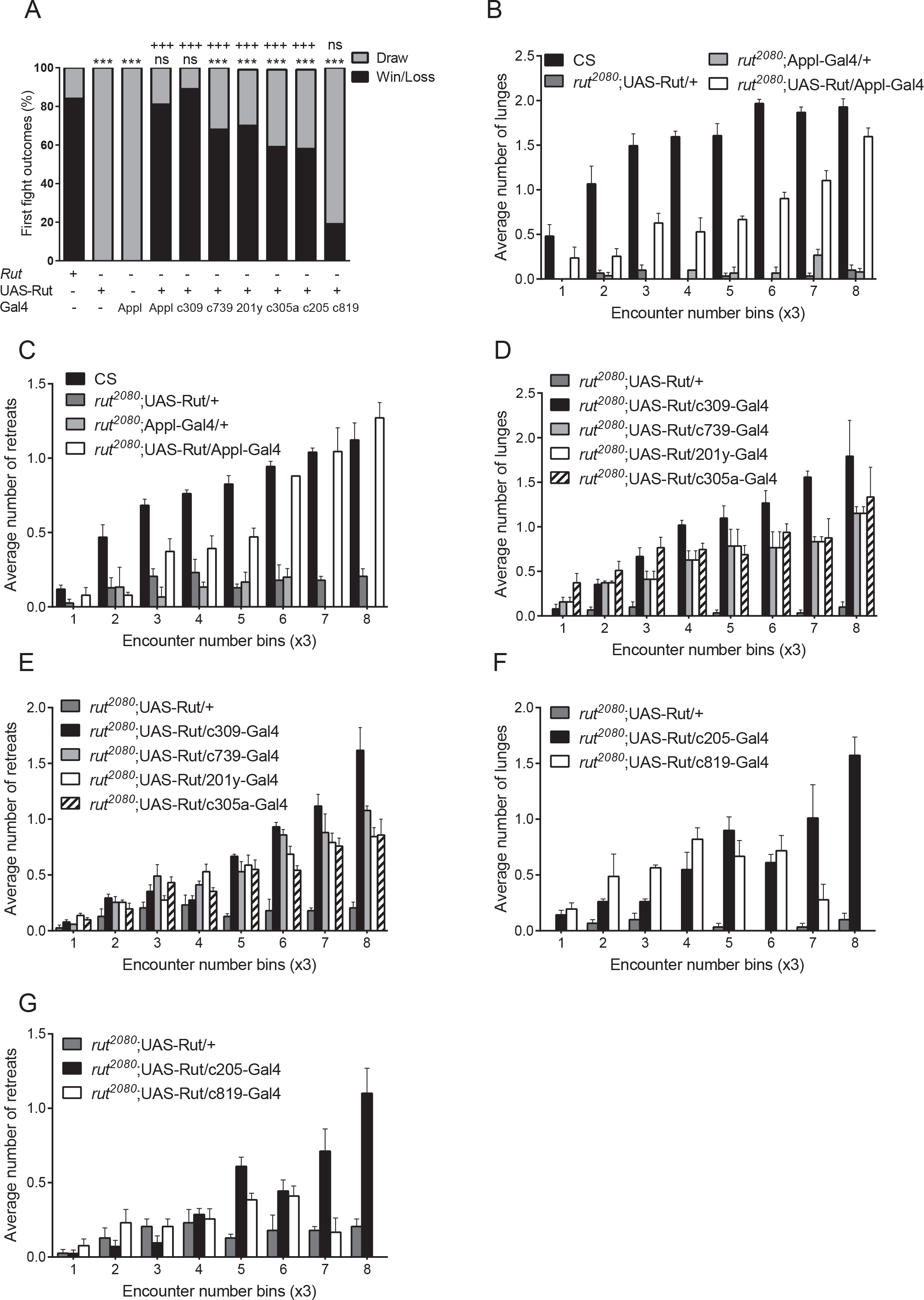
Distinct neuronal circuits mediate behavioral plasticity during agnostic encounters. Restricted expression of *Rut* in specific sub-population of neurons in *rut^2080^* flies yields differential rescue of learning impairment in first fights. (*A*) Lack of development of hierarchies in *rut^2080^* flies is rescued upon expression of UAS-Rut (in a *rut^2080^* background) pan-neuronally (Appl), in the α/β and γ lobes of the MB (c309), α/β lobes of the MB (c739), γ lobes of the MB (201Y), α’/β’ lobes of the MB (c305a) and in the F5 neurons of the FB (c205). No rescue was seen upon reintroduction of *Rut* to the EB (c819). (Two-tailed Chi-square test; ** *P* < 0.01, *** *P* < 0.001, ^+++^ *P* < 0.001, n≥15). *, indicates comparison with CS and +, with *rut^2080^;* UAS-Rut/+. (*B*)-(*G*) Ability to progressively increase lunges/retreats was rescued (relative to *rut^2080^* flies) by expressing *Rut* either pan-neuronally (Appl; *P* < 0.001 for lunges and retreats) or in α/β and γ (c309; *P* < 0.001 for lunges and retreats), α/β alone (c739; *P* < 0.001 for lunges, *P* < 0.01 for retreats), α’/β’ (c305a; *P* < 0.01 for lunges, *P* < 0.05 for retreats) alone, γ (201Y; *P* < 0.001 for lunges, *P* < 0.05 for retreats) alone and in the FB (c205; *P* < 0.001 for lunges, *P* < 0.01 for retreats). EB (c819; *P* > 0.05 for lunges and retreats) expression did not show any rescue. (*B*) and (*C*) are comparisons of Appl-Gal4 with CS and no Gal4 control, (*D*) and (*E*) are comparisons of MB restricted Gal4 drivers (c309, c739, 201y and c305a) with no Gal4 control and (*F*) and (*G*) comparisons are of central complex restricted Gal4 drivers (c205 and c819) with no Gal4 control (Two factor repeated measures ANOVA followed by post hoc Tukey’s multiple comparison test, ‘*P*’ is the interaction term, mean ± SEM, n≥15). The data set for control CS flies, which were trained at the same time of day as experimental lines, is the same as shown in Fig 1. Total number of lunges/retreats in every three successive encounters is analyzed and their mean is reported. SEM: Standard error of the mean

**Table I.**
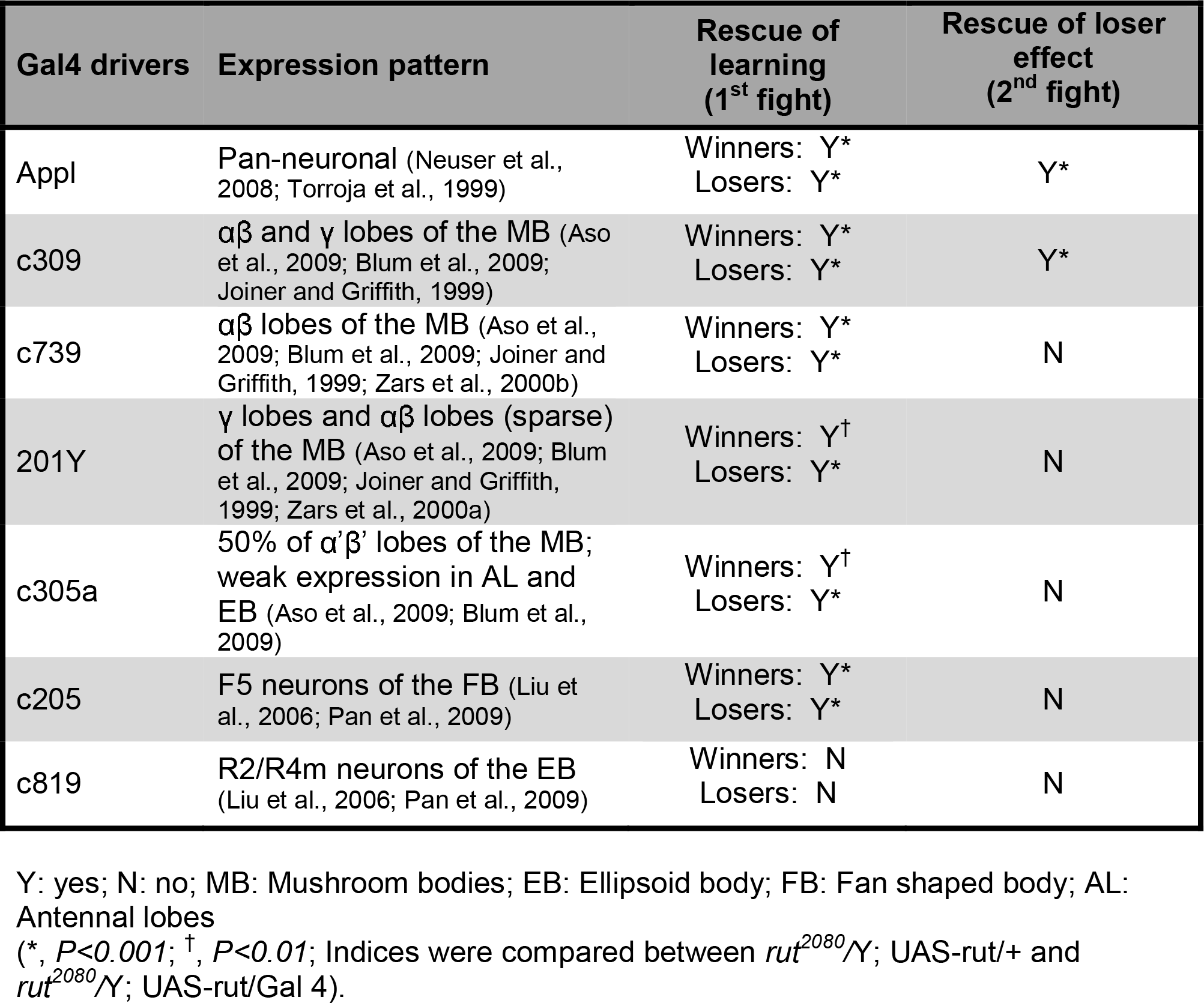
Expression patterns of Gal4 drivers used and rescue results

Analysis of behavioral patterns revealed that distinct neuronal circuits mediate *rut*- dependent behavioral plasticity within a fight (Figure 4 and Table 1). Pan-neuronal expression of *Rut* using the *App*/-Gal4 driver rescued the inability to alter the frequency of use of lunges and retreats seen in *rut^2080^* mutants (Figure 4B, C; *P* < 0.01). We also observed a complete rescue of behavioral transitions pertaining lunges and retreats with *Rut* expression limited to the α/β and γ lobe neurons of the mushroom body using the c309 driver (Figure 4D, E; *P* < 0.001). Furthermore, expression of *Rut*, independently in the α/β, α’/β’ or γ neurons of the MB or in the FB also rescued the deficit in the progressive increase of lunges and retreats seen in *rut^2080^* flies (Figure 4D, E, F, G; *P* < 0.01). The ellipsoid body (EB) of the central complex does not appear to be involved in this function as no rescue was observed with the EB only driver (Figure 4F, G; *P* > 0.05). Restricted expression of *Rut* in specific brain regions rescued the behavioral plasticity of lunges/retreats and the ability to establish dominance hierarchies within a fight.

To confirm that experience-dependent behavioral flexibility was a major substrate of the rescue and not an indirect effect of changes in other general features of aggression, we evaluated multiple aggression related parameters. As the rescue was strongest (comparable to wild-type) with the *Appl*-Gal4 (pan neuronal) and c309 (α/β and γ neurons of the MB) drivers, these were used to evaluate changes in aggressiveness. However, no change in the frequency of encounters, aggression vigor and latencies to engage was found (Figure 5A–E; *P* > 0.05). These results suggest that though pan neuronal and α/β and γ drivers fully rescue the ability to alter the intensities of specific fighting patterns during a fight and the establishment of dominance hierarchies, they do not affect the levels of aggression in these animals.

**Figure 5.**
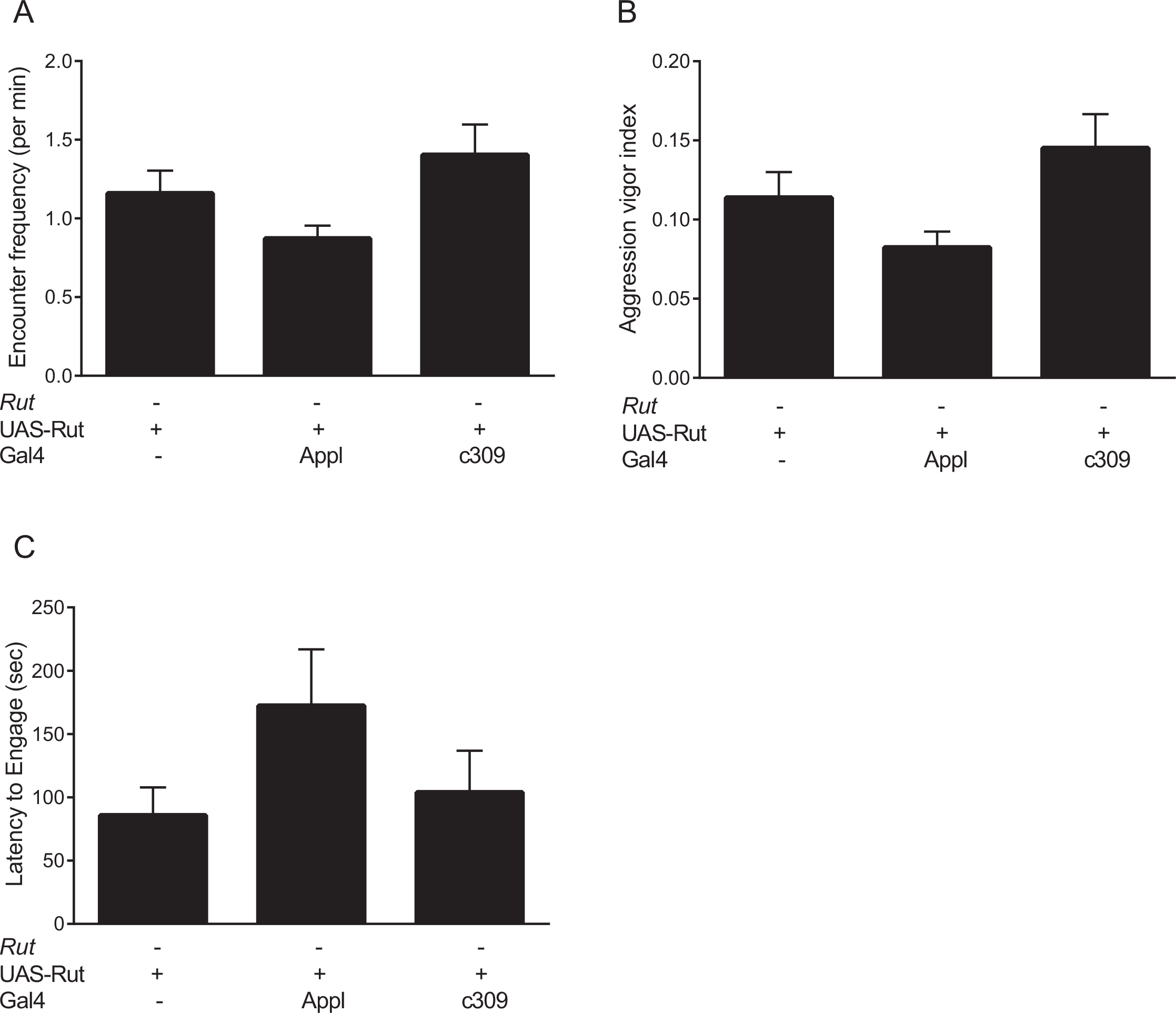
Aggression levels are unchanged upon rescue of aggression-associated learning. (*A*) In comparison to *rut^2080^*;UAS-Rut/+ control flies, *Rut* expression in α/β and γ lobes of the MB (c309; *P* > 0.05) or pan-neuronally (Appl; *P* > 0.05) resulted in a comparable number of encounter frequencies (One factor ANOVA followed by post hoc Dunnett’s multiple comparison test, mean ± SEM, n=20). (B) There is no significant improvement, as compared to *rut^2080^*;UAS-Rut/+ control flies, in aggression vigor index in response to *Rut* expression either pan-neuronally or in the α/β and γ lobes (*P* > 0.05; One factor ANOVA followed by post hoc Dunnett’s multiple comparison test, mean ± SEM, n=20). (*C*) Latency to engage in an encounter (*P* > 0.05) remains comparable upon *Rut* expression using either Appl or c309 Gal4 driver lines in comparison to *rut^2080^*;UAS- Rut/+ control flies (One factor ANOVA followed by post hoc Dunnett’s multiple comparison test, mean ± SEM, n=20). SEM: Standard error of the mean.

Our analysis indicates that in accordance with the multimodal inputs associated with dyadic interactions, multiple neuronal groups in the MB (α/β, α’/β’, γ) and CC (FB) are involved in the processing of these inputs that lead to the progressive changes in offensive and defensive strategies within a fight. The rescue experiments also highlight the role of cAMP signaling in behavioral plasticity in aggression to be independent of general changes in aggression levels.

### cAMP signaling is necessary for the development of the loser effect

To test if aggression-associated hierarchies developed in the first fights influenced the outcomes of subsequent fights, losers from first fights were paired with unfamiliar, socially naive opponents, after a 60-minute rest period. Dominance relationships were assigned using the three lunge- three retreat rule as in the first fights. Fights that fell under the ‘draw’ category were further subdivided into three groups. ‘High intensity’ draws consisted of experienced flies predominantly using lunges in their second fight without satisfying 3 lunge-3 retreat criteria for dominance relationships. Similarly, ‘low intensity’ draws consisted of experienced flies predominantly using retreats. Fights in which the flies did not engage in agonistic interactions were considered as ‘no intensity’ and categorized as draws. For statistical analysis, high and low intensity draws were grouped with wins and losses, respectively. Consistent with previous studies (Trannoy et al., 2016; Yurkovic et al., 2006), wild-type CS loser flies always lost to naïve opponents in their second fights (Figure 6A). This indicates that an experience of social loss results in a strong loser effect in subsequent fights.

**Figure 6.**
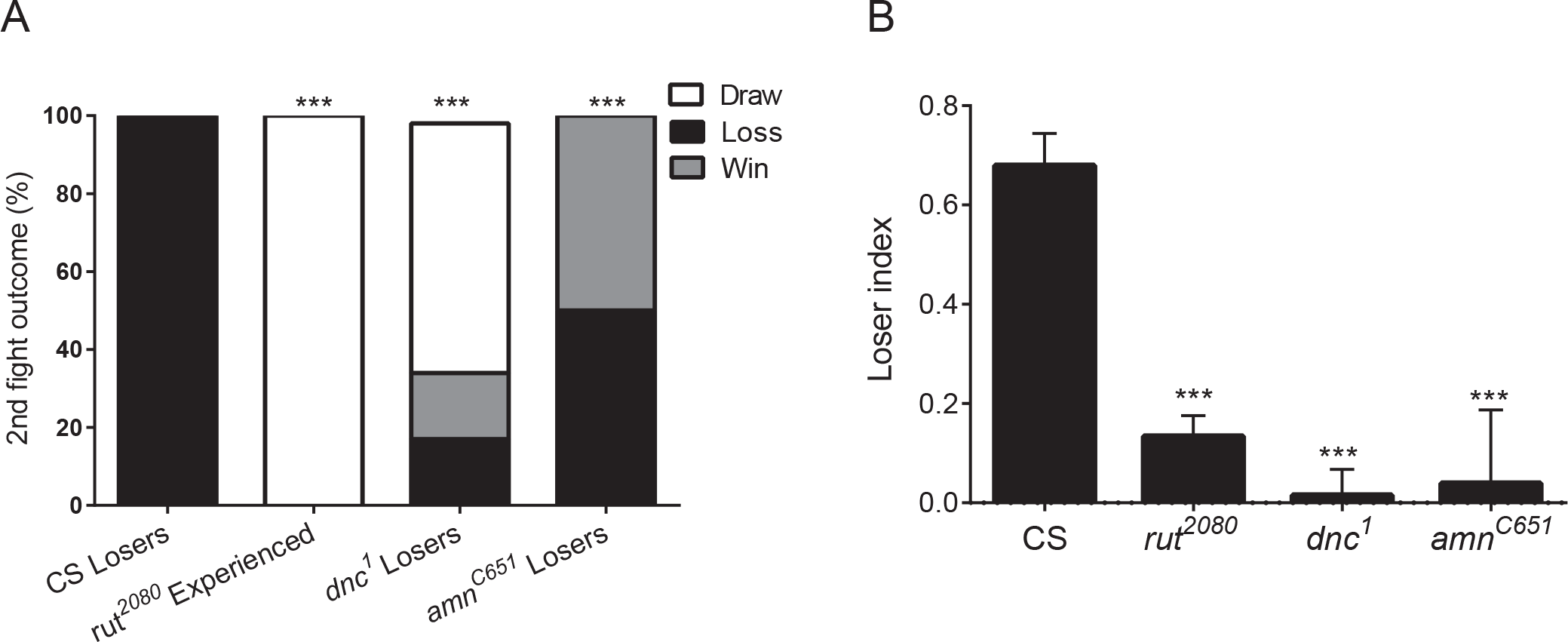
cAMP signaling is necessary for developing loser mentality. (*A*) CS losers lost all their second fights against naive opponents while *amn^C651^* flies lost and won equal number of fights. *dnc^1^* flies show weak dominance relationships and their interactions largely ended in draws, again suggestive of memory defects. All previously experienced *rut^2080^* flies engaged in draws. (Two-tailed Chi-square test; *** *P* < 0.001, n≥20). *, indicates comparison with CS. (*B*) Loser index assesses the experience dependent modulation of fighting strategies of experienced flies in their second fights against a naive opponent. *rut^2080^, dnc^1^* and *amn^C651^* flies, all showed severely compromised loser indices compared to CS. (One factor ANOVA followed by post hoc Dunnett’s multiple comparison test, *** *P* < 0.001, mean ± SEM, n≥20). *, indicates comparison with CS.

As *rut^2080^* flies did not generate winners or losers in the first fight, both the individuals were considered as experienced and used in second fights. However, as in their first fights, *rut^2080^* mutants were unable to form any dominance relationships in their second fights (Figure 6A; *P* < 0.001). *dnc^1^* also displayed no significant loser effect with 18% losses, 18% wins and 64% draws (Figure 6A; *P* < 0.001).

Surprisingly, *amn^C651^* flies did not show any loser effect. 50% of analyzed second fights resulted in losses and the remaining in wins for losers (Figure 6A; *P* < 0.001). Unlike *rut* and *dnc, amn^C651^* flies develop dominant-subordinate relations as efficiently as CS in their first fights (Figure 1C) but are still unable to consolidate this experience into a loser effect. This suggests a special requirement of Amn gene product in stabilizing the hierarchical structures in flies in the form of a loser effect.

A loser index, calculated as the difference between the number of encounters lost and the number of encounters won divided by the total number of encounters in the second fight (see Materials and Methods), was used to represent the loser effect as a consequence of past experience. This allowed direct evaluation of experience-dependent alterations in fighting strategy in the losers across genotypes in subsequent fights. CS flies showed a high loser index consistent with their robust loser effect (Figure 6B). In contrast, the *amn^C651^* and *dnc^1^* losers, and the experienced *rut^2080^* flies displayed significantly lower loser indices (Figure 6B; *P* < 0.001).

These results show that *dnc^1^* and *rut^2080^* flies, which display compromised development of dominance hierarchies in their first fights, did not display a loser effect. In contrast, *amn^C651^*, which could form dominance hierarchies in their first fights, also lacked the loser effect. *Amn* appears to have an independent function from *Rut* and *Dnc* and is necessary for the stabilization of the social status from the first fight in the form of a loser effect.

### α/β and γ lobes of the MB cooperate to mediate cAMP-dependent loser effect

Previous experience of loss results in a robust loser effect in wild-type flies. We investigated the neurogenetic circuitry involved in the development of this effect.

In the second fights, both *Appl* and c309 Gal4 driver lines were able to restore the loser effect in *rut^2080^* flies to levels statistically indistinguishable from wild-type CS flies (Figure 7A, B and Table 1). Pan neuronal (Appl) or combined α/β and γ lobe-specific (c309) expression resulted in most losers losing and a small proportion of non-aggressive draws (Figure 7A; *P* < 0.001). Loser index in these flies was significantly greater than controls (Figure 7B; *P* < 0.001) and comparable to wild-type flies (Figure 7B; *P* > 0.05).

**Figure 7.**
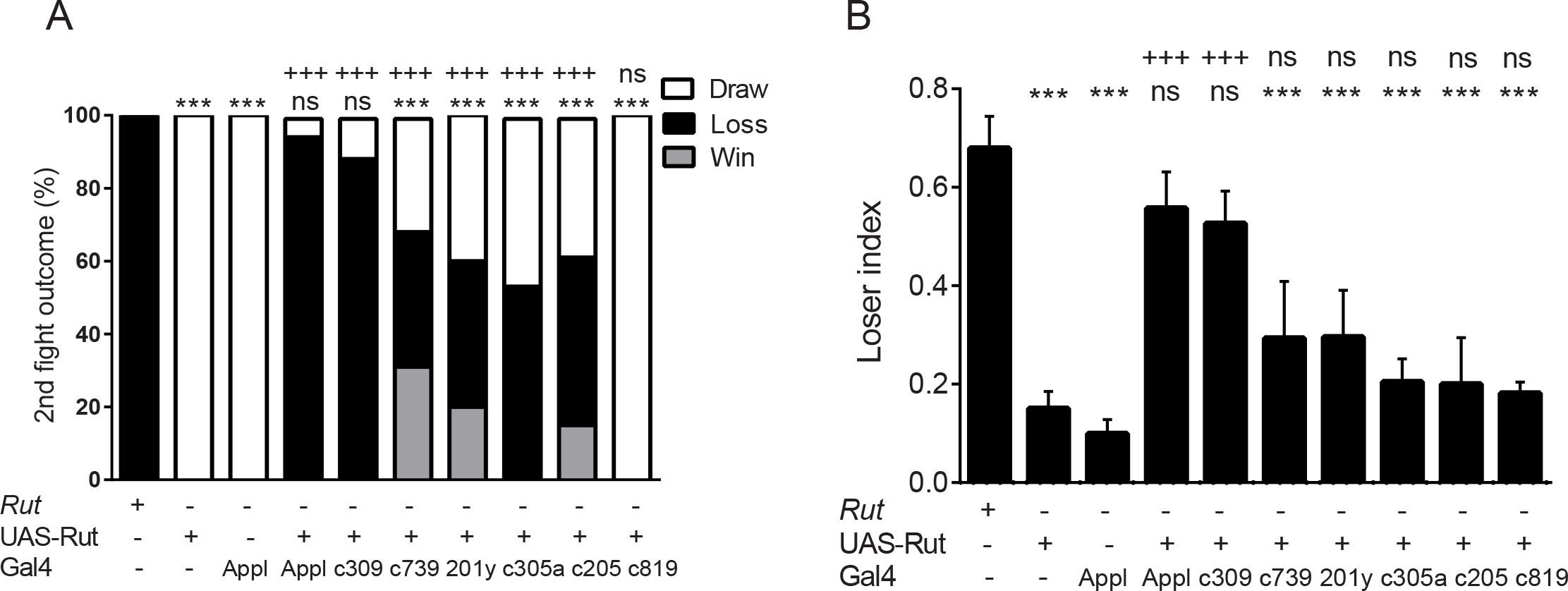
Recruitment of specific neuronal circuits for the development of the loser effect. (*A*) Rescue experiments expressing *Rut* (in a *rut^2080^* background) either pan-neuronally (Appl) or in the α/β and γ lobes of the MB (c309) showed a strong recovery of the loser effect. Expression in the α/β lobes (c739), γ lobes (201Y), α’/β’ lobes (c305a) and in the FB (c205) of *rut* mutants resulted in partial recovery of dominance relationships. No rescue was seen upon the reintroduction of *Rut* in the EB (c819) of *rut^2080^*. (Two-tailed Chi-square test;*** *P* < 0.001, +++ *P* < 0.001 n≥15). *, indicates comparison with CS and +, with *rut^2080^*; UAS-Rut/+. (*B*) Loser index analysis revealed that pan-neuronal and α/β and γ lobe combined expression is able to rescue the loser effect. All other combinations did not show any rescue. (One factor ANOVA followed by post hoc Tukey’s multiple comparison test, *** *P* < 0.001, +++ *P* < 0.001, mean ± SEM, n≥15). *, indicates comparison with CS and +, with *rut^2080^;* UAS-Rut/+. SEM: Standard error of the mean. The data set for control CS flies, which were trained at the same time of day as experimental lines, is the same as shown in Fig 6.

Interestingly, expression of UAS-Rut limited to individual substructures of the MB and CC was not sufficient to rescue impaired loser effect observed in *rut* flies (Figure 7B; Table 1). Expression in individual neuronal subpopulations using the c739, 201Y, c305a or c205 Gal4 drivers (in the *rut^2080^* background), resulted in a higher proportion of non-draw outcomes in losers paired with naïve opponents but failed to show a robust loser effect (Figure 7A; *P* < 0.001). As expected, there was no significant change in the loser index for these flies (Figure 7B; *P* > 0.05). Consistent with compromised learning in the first fights, expression limited to the EB using the c819 Gal4 driver did not rescue the loser effect (Figure 7; *P* > 0.05).

Our results implicate combined processing by the α/β and γ neurons of the MBs in the development of the loser effect.

## DISCUSSION

Two distinct experience-dependent behavioral modifications are observed in *Drosophila* aggression. The first results in dominance relationships correlated with real time, progressive changes in fighting strategies. The second is the consolidation of the experience of a previous loss to a loser effect in subsequent fights. Our data indicate that these behavioral changes involve specific cAMP-mediated signaling and are processed by overlapping neuronal circuits involving two temporally separated memory traces.

Rut and Dnc enzymes have been implicated in learning and/or STM in multiple paradigms, including classical conditioning and courtship-associated learning and memory (Bragina and Kamyshev, 2003; Gailey et al., 1984; Livingstone et al., 1984; McGuire et al., 2005). *rut* and *dnc* flies do not develop dominance hierarchies as well as CS flies, with all *rut-rut* fights resulting in draws. Compared to CS, both *rut* and *dnc* are unable to display experience-dependent modification of fighting strategies and are deficient in generating losers and winners.

*rut* and *dnc* mutant flies are deficient in memory formation in various behavioral assays (Dudai et al., 1976; Gailey et al., 1984; Livingstone et al., 1984; McGuire et al., 2005). In olfactory conditioning, both *rut* and *dnc* mutants demonstrate rapid memory degradation within an hour of training but have no impact on later memory (Chen et al., 1986; Levin et al., 1992; Tamura et al., 2003). Also, *rut* mutant flies were unable to demonstrate robust 30 min-memory after an hour of training in a courtship conditioning paradigm (Ishimoto et al., 2013). Therefore, *rut* and *dnc* mutants’ inability to alter behavioral states might be due to poor memory acquisition pertaining winner-loser status, which results in lack of hierarchical relationships in a fight.

The Amn gene product encodes for three putative neuropeptides (Feany and Quinn, 1995 and Moore et al., 1998). Genetic and biochemical studies have implicated the Amn locus in cAMP dynamics (Feany and Quinn, 1995; Moore et al., 1998; Bhattacharya et al., 2004). It is assumed, from analogous data in other organisms, that the Amn gene product/s activate adenylyl cyclase activity via GPCR signaling (Abel and Kandel, 1995; McGuire et al., 2005). However, there is no direct evidence of the Amn gene products being secreted and acting as a classical neuropeptide via GPCR activation. In our experiments, *amn* flies are as competent in modifying fighting strategies and establishing hierarchal relationships as CS flies but are unable to retain the loser status in subsequent fights. These results suggest that the plasticity in modifying aggressive strategies during a fight is mediated by cAMP dynamics under the control of Rut and Dnc gene products (but not Amn) and is necessary to establish dominance structures.

All three cAMP pathway mutants displayed a reduction in aggression vigor as compared to CS. This includes *amn*, which displays normal behavioral flexibility and dominance hierarchies. These results suggest that the level of aggression in flies is independent of their ability to modify behavior in an experience-dependent manner. The latency to engage in fights in cAMP mutants was also comparable to CS. The latter suggests no overt changes in motivational states and support the previous conclusion.

Brain region specific expression of the Rut gene product in a *rut* mutant background implicates specific neuronal circuits in status-dependent behavioral changes during agonistic encounters. All the drivers that rescue the establishment of dominance also restore the ability to alter the usage of lunges and retreats during a fight.

Rescue by Appl or c309 did not alter the encounter frequency, aggression vigor or the latency to engage displayed by *rut* mutants. In line with our analysis of cAMP pathway mutants, these results demonstrate that the rescue of dominance is attributable to a restoration of experience-dependent plasticity mediated by *Rut* in specific circuits and not a consequence of modified aggressiveness.

Our study suggests a functional role for MBs and Central Complex (CC) in learning and memory associated with aggression, consistent with those described in courtship conditioning (Joiner and Griffith, 1999; McBride et al., 1999; Sitnik et al., 2003). Mushroom bodies, which are central to olfactory learning, have been previously correlated with changes in agonistic behavior. *Neura/ized* mutants with altered MB organization have been reported to increase aggressiveness when food is limiting (Rollmann et al., 2008). Inhibition of synaptic output from the MB has also been shown to reduce levels of aggression (Liu et al., 2011).

Interestingly, not only neuronal groups of the MB (the α/β, γ and α’/β’ neurons) were implicated, the F5 neurons of the FB but not the EB of the CC was also found to be functionally involved. The FB has been previously reported to mediate visual learning of specific pattern features (Liu et al., 2006; Pan et al., 2009). A recent study has underscored the importance of the CC in aggression by demonstrating the modulation of aggressive behavior by dopaminergic PPM3 neurons that synapse onto the FB (Alekseyenko et al., 2013). While there is no evidence of direct anatomical connectivity between the MB and the F5 neurons of the FB (Li et al., 2009; Young and Armstrong, 2010), it is possible that they are functionally linked by neuromodulatory dopaminergic signaling.

In the courtship suppression paradigm, multiple biochemically and anatomically distinct neural circuits have been implicated (Joiner and Griffith, 2000). Our observations in aggression-associated behaviors are analogous and possibly reflect the integration of multiple sensory inputs contributing to the establishment of dominance hierarchies.

The reduced aggression seen in cAMP pathway mutants could be due to alteration in the production of aggression-associated signal/s and/or their perception. While this possibility awaits systematic investigation, the rescue of behavioral flexibility without altering aggressiveness seen in our experiments suggests that the two may be separable. It is also possible that the observed behavioral deficits originate from neurodevelopmental functions of these genes. Our study does not explicitly rule this out though adult specific expression of Rut is sufficient to rescue learning and memory defects in other behavioral paradigms (Mao et al., 2004; McGuire et al., 2003; Putz and Heisenberg, 2002; Tan et al., 2010). Future studies will need to address this question specifically in the context of aggression.

The robust loser effect seen in wild-type flies is absent in *rut, dnc* and *amn* mutant lines. As in their first fights, all *rut-rut* second fights end in draws. No loser effect was also seen in *dnc* flies. Both *rut* and *dnc* have compromised development of hierarchies suggesting that learning within a fight and establishing dominance relationships are necessary for the development of the loser effect. Interestingly, *amn* flies also lack the loser effect, though they can adjust fighting strategies and develop dominance relationships as well as wild-type flies. The Amn gene product appears to mediate the consolidation of the social status acquired during the first fight into the loser effect. Alternatively, Amn may be necessary for a distinct memory phase resulting in the loser effect. A similar function for Amn has been reported in olfactory conditioning experiments where *amn* mutants fail to develop intermediate-term memory or are unable to consolidate STM into this relatively longer lasting phase (Feany and Quinn, 1995; Yu et al., 2006). Amn has also been reported to mediate memory stability in courtship conditioning (Siegel and Hall, 1979). Our findings support this model as *amn* flies do establish winner-loser relationships but fail to consolidate these dominance hierarchies. If the Amn gene product functions as a secreted neuropeptide, future rescue experiments involving the cognate receptor/s will be necessary to determine the neural circuits subserving Amn-dependent establishment of the loser effect. A less robust, short duration winner effect has been recently reported in flies (Trannoy and Kravitz, 2016; Trannoy et al., 2016). It remains to be seen if cAMP signaling and circuit features analogous to our observations mediate this phenomenon.

Rescue of *rut* simultaneously in the α/β and γ lobe neurons of the MB, not only fully rescues its ability to generate dominance relationships but also the loser effect. However, *rut* expression independently in the α/β or the γ neurons were unable to restore the loser effect, though social hierarchy was rescued. We hypothesize that two distinct Rut dependent memory traces facilitate the formation of the temporally distinct memory phases. A short-lived engram in distinct MB and CC substructures mediates behavioral plasticity within a fight leading to the formation of dominance relationships. Combinatorial processing between α/β and γ lobe neurons of the MB enable formation of a second, longer lasting memory trace that is necessary for the establishment of the loser effect.

Olfactory conditioning studies have suggested that the α/β neurons are indispensable for memory consolidation and retrieval, while odor-shock coincidence detection maps to the y neurons (Dubnau and Chiang, 2013; Dubnau et al., 2001; Qin et al., 2012). These observations demonstrate recruitment of specific subsets of MB neurons for initial associations and others for memory consolidation suggesting circuit level coordination between MB substructures. Our experiments provide evidence towards such systems level memory consolidation in aggression where an association between α/β and γ lobes of the MB is critical for the establishment of the loser effect. However, the existence of two parallel, independent traces with differing kinetics of formation and decay cannot be formally ruled out.

Experience-dependent modification of innate behaviors involves multiple components and previous studies have demonstrated the central importance of pheromonal, aminergic and other modulatory activities in aggressive behavior (Alekseyenko et al., 2014; Alekseyenko et al., 2013; Andrews et al., 2014; Certel et al., 2010; Chan and Kravitz, 2007; Dankert et al., 2009; Dierick and Greenspan, 2007; Hoopfer, 2016; Hoyer et al., 2008; Kohl et al., 2015; Liu et al., 2011; Luo et al., 2014; Wang and Anderson, 2010; Yuan et al., 2014; Zwarts et al., 2012). However, these studies focus on aggression levels and do not directly assess behavioral plasticity and memory components in aggression. A recent study indicated that *rut^2080^* and *amn^1^* are unable to demonstrate dominance hierarchies due to compromised aggression (Trannoy et al., 2016). However, this study did not explore the lack of behavioral flexibility in these mutants. In our study, *rut^2080^* has low proclivity to engage in the first 10 min of fights, fail to form dominance relations but are capable of executing high intensity maneuvers. In contrast, *amn^C651^* (a functionally strong allele of *amn* (Rosay et al., 2001)), showed reduced aggression vigor but was still able to establish dominance hierarchies. Further, the rescue of *Rut* restores dominance patterns but not aggressiveness. Therefore, low aggression is not correlated with the inability to establish winner-loser statuses in a fight.

cAMP signaling appears to have a conserved role in establishing dominance relationships across phyla. As observed previously in the olfactory conditioning and courtship paradigms, *rut* and *dnc* mutants display similar behavioral deficits in aggression-associated behavioral flexibility despite having opposing biochemical functions. Amn activity is also thought to increase cAMP levels (perhaps with different kinetics than Rut). Thus, cAMP levels and fine-tuning of cAMP levels may encode signal integration by the cAMP second messenger system. In crayfish, accumulation of cAMP is necessary to form the loser effect while a reduction in cAMP levels is associated with the winner effect (Momohara et al., 2016). Biogenic amines have been associated with aggression and winner/loser effects in animals including flies (Alekseyenko et al., 2014; Alekseyenko et al., 2013; Andrews et al., 2014; Hoyer et al., 2008; Zhou and Rao, 2008), crayfish (Momohara et al., 2016; Panksepp, 2003), crickets (Rillich and Stevenson, 2014; Rillich and Stevenson, 2015; Stevenson et al., 2005) and mammals (de Almeida et al., 2005). Aminergic signaling commonly impinges on cAMP/Protein Kinase A (PKA) pathways implicating the cAMP signaling as a central effector of neuromodulation by biogenic amines (Neckameyer and Leal, 2017). In mammals, the cAMP pathway has been shown to influence aggressive behavior (Breuillaud et al., 2012). cAMP signaling in the basolateral amygdala has also been correlated with memory associated with conditioned defeat, a paradigm analogous to the loser effect (Jasnow et al., 2005; Markham et al., 2010). At the circuit level, recent work in zebrafish has implicated antagonistic regulation by two sub-regions of the dorsal habenula to establish winner-loser status (Chou et al., 2016).

Our results implicate sequential recruitment of cAMP signaling components with Rut and Dnc activities required for behavioral flexibility within a fight and, consequently, to establish dominance hierarchies. Behavioral plasticity, rather than aggressiveness, is correlated with social status. Establishment of social status facilitates the formation of a more stable and longer lasting memory phase that requires the additional involvement of the Amn gene product. Neuronal circuits subserving aggression associated learning and memory show phasic recruitment. While a short-lived Rut-dependent trace in multiple MB and CC substructures mediate learning during agonistic encounters, combinatorial processing by both α/β and γ lobe neurons of the MB, is necessary for the development of the longer lasting loser effect.

This study provides mechanistic insight into circuit level associations specific to different phases of behavioral plasticity and memory in *Drosophila* aggression and the integration of biochemical signaling at the single neuron level with systems level consolidation.

## ACKNOWLEDGEMENTS

We thank Dr D. Barua (IISER Pune, India) for advice on statistical analysis. We are grateful to Prof. R. Strauss (University of Mainz, Germany) for providing Canton S, *rut^2080^* and *amn^c651^* fly lines. Prof. N. K. Subhedar (IISER Pune, India), Prof. L. S. Shashidhara (IISER Pune, India), Dr. M. Lahiri (IISER Pune, India), Dr. R. Rajan (IISER Pune, India) and Prof. K. S. Krishnan (NCBS Bangalore, India) are acknowledged for discussions and critical reading of earlier versions of the manuscript.

## REFERENCES

Alekseyenko, O. V., Chan, Y. B., Fernandez, M. P., Bulow, T., Pankratz, M. J. and Kravitz, E. A. (2014). Single serotonergic neurons that modulate aggression in Drosophila. Curr Biol 24, 2700–7.

Alekseyenko, O. V., Chan, Y. B., Li, R. and Kravitz, E. A. (2013). Single dopaminergic neurons that modulate aggression in Drosophila. Proc Natl Acad Sci USA 110, 6151–6.

Ali, Y., Escala, W., Ruan, K. and Zhai, R. (2011). Assaying locomotor, learning, and memory deficits in Drosophila models of neurodegeneration. J Vis Exp, e2504.

Andrews, J. C., Fernandez, M. P., Yu, Q., Leary, G. P., Leung, A. K., Kavanaugh, M. P., Kravitz, E. A. and Certel, S. J. (2014). Octopamine neuromodulation regulates Gr32a-linked aggression and courtship pathways in Drosophila males. PLoS Genet 10, e1004356.

Asahina, K., Watanabe, K., Duistermars, B. J., Hoopfer, E., Gonzalez, C. R., Eyjolfsdottir, E. A., Perona, P. and Anderson, D. J. (2014). Tachykinin-expressing neurons control male-specific aggressive arousal in Drosophila. Cell 156, 221–35.

Aso, Y., Grubel, K., Busch, S., Friedrich, A. B., Siwanowicz, I. and Tanimoto, H. (2009). The mushroom body of adult Drosophila characterized by GAL4 drivers. J Neurogenet 23, 156–72.

Blum, A. L., Li, W., Cressy, M. and Dubnau, J. (2009). Short- and long-term memory in Drosophila require cAMP signaling in distinct neuron types. Curr Biol 19, 1341–50.

Bragina, Y. V. and Kamyshev, N. G. (2003). Comparative studies of four Drosophila P-insertion mutants with memory defects. Neurosci Behav Physiol 33, 73–9.

Brembs, B. (2003). Operant conditioning in invertebrates. Curr Opin Neurobiol 13, 710–7.

Breuillaud, L., Rossetti, C., Meylan, E. M., Merinat, C., Halfon, O., Magistretti, P. J. and Cardinaux, J. R. (2012). Deletion of CREB-regulated transcription coactivator 1 induces pathological aggression, depression-related behaviors, and neuroplasticity genes dysregulation in mice. Biol Psychiatry 72, 528–36.

Busto, G., Cervantes-Sandoval, I. and Davis, R. (2010). Olfactory learning in Drosophila. Physiology 25, 338–346.

Certel, S. J., Leung, A., Lin, C. Y., Perez, P., Chiang, A. S. and Kravitz, E. A. (2010). Octopamine neuromodulatory effects on a social behavior decision-making network in Drosophila males. PLoS One 5, e13248.

Chan, Y.-B. and Kravitz, E. (2007). Specific subgroups of FruM neurons control sexually dimorphic patterns of aggression in Drosophila melanogaster. Proc Natl Acad Sci USA 104, 19577–19582.

Chen, C. N., Denome, S. and Davis, R. L. (1986). Molecular analysis of cDNA clones and the corresponding genomic coding sequences of the Drosophila dunce+ gene, the structural gene for cAMP phosphodiesterase. Proc Natl Acad Sci U S A 83, 9313–7.

Chen, S., Lee, A. Y., Bowens, N. M., Huber, R. and Kravitz, E. A. (2002). Fighting fruit flies: a model system for the study of aggression. Proc Natl Acad Sci USA 99, 5664–8.

Chou, M. Y., Amo, R., Kinoshita, M., Cherng, B. W., Shimazaki, H., Agetsuma, M., Shiraki, T., Aoki, T., Takahoko, M., Yamazaki, M. et al. (2016). Social conflict resolution regulated by two dorsal habenular subregions in zebrafish. Science 352, 87–90.

Dankert, H., Wang, L., Hoopfer, E., Anderson, D. and Perona, P. (2009). Automated monitoring and analysis of social behavior in Drosophila. Nat Meth 6, 297303.

Davis, R. L. and Kiger, J. A. (1981). Dunce mutants of Drosophila melanogaster. mutants defective in the cyclic AMP phosphodiesterase enzyme system. J Cell Biol 90, 101–7.

de Almeida, R. M., Ferrari, P. F., Parmigiani, S. and Miczek, K. A. (2005). Escalated aggressive behavior: dopamine, serotonin and GABA. Eur J Pharmacol 526, 51–64.

Dierick, H. A. and Greenspan, R. J. (2006). Molecular analysis of flies selected for aggressive behavior. Nat Genet 38, 1023–31.

Dierick, H. A. and Greenspan, R. J. (2007). Serotonin and neuropeptide F have opposite modulatory effects on fly aggression. Nat Genet 39, 678–82.

Dubnau, J. and Chiang, A. S. (2013). Systems memory consolidation in Drosophila. Curr Opin Neurobiol 23, 84–91.

Dubnau, J., Grady, L., Kitamoto, T. and Tully, T. (2001). Disruption of neurotransmission in Drosophila mushroom body blocks retrieval but not acquisition of memory. Nature 411, 476–80.

Dudai, Y., Corfas, G. and Hazvi, S. (1988). What is the possible contribution of Ca^2^+-stimulated adenylate cyclase to acquisition, consolidation and retention of an associative olfactory memory in Drosophila. J Comp Physiol A 162, 101–9.

Dudai, Y., Jan, Y. N., Byers, D., Quinn, W. G. and Benzer, S. (1976). dunce, a mutant of Drosophila deficient in learning. Proc Natl Acad Sci USA 73, 1684–8.

Edwards, A., Zwarts, L., Yamamoto, A., Callaerts, P. and Mackay, T. (2009). Mutations in many genes affect aggressive behavior in Drosophila melanogaster. BMC biology 7, 29.

Feany, M. B. and Quinn, W. G. (1995). A neuropeptide gene defined by the Drosophila memory mutant amnesiac. Science 268, 869–73.

Gailey, D., Jackson, F. and Siegel, R. (1984). Conditioning mutations in Drosophila melanogaster affect an experience dependent behavioral modification in courting males. Genetics 106, 613–623.

Hoopfer, E. D. (2016). Neural control of aggression in Drosophila. Curr Opin Neurobiol 38, 109–18.

Hoopfer, E. D., Jung, Y., Inagaki, H. K., Rubin, G. M. and Anderson, D. J. (2015). P1 interneurons promote a persistent internal state that enhances inter-male aggression in Drosophila. Elife 4.

Hoyer, S. C., Eckart, A., Herrel, A., Zars, T., Fischer, S. A., Hardie, S. L. and Heisenberg, M. (2008). Octopamine in male aggression of Drosophila. Curr Biol 18, 159–67.

Ishimoto, H., Wang, Z., Rao, Y., Wu, C. F. and Kitamoto, T. (2013). A novel role for ecdysone in Drosophila conditioned behavior: linking GPCR-mediated non- canonical steroid action to cAMP signaling in the adult brain. PLoS Genet 9, e1003843.

Jasnow, A. M., Shi, C., Israel, J. E., Davis, M. and Huhman, K. L. (2005). Memory of social defeat is facilitated by cAMP response element-binding protein overexpression in the amygdala. Behav Neurosci 119, 1125–30.

Joiner, M. and Griffith, L. (1999). Mapping of the anatomical circuit of CaM kinase-dependent courtship conditioning in Drosophila. Learn Mem 6, 177–192.

Joiner, M. A. and Griffith, L. C. (2000). Visual input regulates circuit configuration in courtship conditioning of Drosophila melanogaster. Learn Mem 7, 32–42.

Kamyshev, N. G., Smirnova, G. P., Kamysheva, E. A., Nikiforov, O. N., Parafenyuk, I. V. and Ponomarenko, V. V. (2002). Plasticity of social behavior in Drosophila. Neurosci Behav Physiol 32, 401–8.

Kohl, J., Huoviala, P. and Jefferis, G. S. (2015). Pheromone processing in Drosophila. Curr Opin Neurobiol 34, 149–57.

Levin, L. R., Han, P. L., Hwang, P. M., Feinstein, P. G., Davis, R. L. and Reed, R. R. (1992). The Drosophila learning and memory gene rutabaga encodes a Ca^2+^/Calmodulin-responsive adenylyl cyclase. Cell 68, 479–89.

Lim, R. S., Eyjolfsdottir, E., Shin, E., Perona, P. and Anderson, D. J. (2014). How food controls aggression in Drosophila. PLoS One 9, e105626.

Liu, G., Seiler, H., Wen, A., Zars, T., Ito, K., Wolf, R., Heisenberg, M. and Liu, L. (2006). Distinct memory traces for two visual features in the Drosophila brain. Nature 439, 551–6.

Liu, W., Liang, X., Gong, J., Yang, Z., Zhang, Y. H., Zhang, J. X. and Rao, Y. (2011). Social regulation of aggression by pheromonal activation of Or65a olfactory neurons in Drosophila. Nat Neurosci 14, 896–902.

Livingstone, M. S., Sziber, P. P. and Quinn, W. G. (1984). Loss of calcium/calmodulin responsiveness in adenylate cyclase of rutabaga, a Drosophila learning mutant. Cell 37, 205–15.

Luo, J., Lushchak, O. V., Goergen, P., Williams, M. J. and Nassel, D. R. (2014). Drosophila insulin-producing cells are differentially modulated by serotonin and octopamine receptors and affect social behavior. PLoS One 9, e99732.

Mao, Z., Roman, G., Zong, L. and Davis, R. L. (2004). Pharmacogenetic rescue in time and space of the rutabaga memory impairment by using Gene-Switch. Proc Natl Acad Sci U S A 101, 198–203.

Markham, C. M., Taylor, S. L. and Huhman, K. L. (2010). Role of amygdala and hippocampus in the neural circuit subserving conditioned defeat in Syrian hamsters. Learn Mem 17, 109–16.

McBride, S., Giuliani, G., Choi, C., Krause, P., Correale, D., Watson, K., Baker, G. and Siwicki, K. (1999). Mushroom body ablation impairs short-term memory and long-term memory of courtship conditioning in Drosophila melanogaster. Neuron 24, 967–977.

McGuire, S. E., Deshazer, M. and Davis, R. L. (2005). Thirty years of olfactory learning and memory research in Drosophila melanogaster. Prog Neurobiol 76, 328–47.

McGuire, S. E., Le, P. T., Osborn, A. J., Matsumoto, K. and Davis, R. L. (2003). Spatiotemporal rescue of memory dysfunction in Drosophila. Science (New York, N.Y.) 302, 1765–1768.

Momohara, Y., Minami, H., Kanai, A. and Nagayama, T. (2016). Role of cAMP signalling in winner and loser effects in crayfish agonistic encounters. Eur J Neurosci 44, 1886–95.

Moore, M. S., DeZazzo, J., Luk, A. Y., Tully, T., Singh, C. M. and Heberlein, U. (1998). Ethanol intoxication in Drosophila: Genetic and pharmacological evidence for regulation by the cAMP signaling pathway. Cell 93, 997–1007.

Neckameyer, W. and Leal, S. (2017). Diverse Functions of Insect Biogenic Amines as Neurotransmitters, Neuromodulators, and Neurohormones. In Hormones, Brain and Behavior, pp. 367–401: Elsevier

Neuser, K., Triphan, T., Mronz, M., Poeck, B. and Strauss, R. (2008). Analysis of a spatial orientation memory in Drosophila. Nature 453, 1244–1247.

Pan, Y., Zhou, Y., Guo, C., Gong, H., Gong, Z. and Liu, L. (2009). Differential roles of the fan-shaped body and the ellipsoid body in Drosophila visual pattern memory. Learn Mem 16, 289–95.

Panksepp, J. (2003). Neuroscience. Feeling the pain of social loss. Science 302, 237–9.

Penn, J. K., Zito, M. F. and Kravitz, E. A. (2010). A single social defeat reduces aggression in a highly aggressive strain of Drosophila. Proc Natl Acad Sci USA 107, 12682–6.

Putz, G. and Heisenberg, M. (2002). Memories in drosophila heat-box learning. Learn Mem 9, 349–359.

Qin, H., Cressy, M., Li, W., Coravos, J. S., Izzi, S. A. and Dubnau, J. (2012). Gamma neurons mediate dopaminergic input during aversive olfactory memory formation in Drosophila. Curr Biol 22, 608–14.

Ramin, M., Domocos, C., Slawaska-Eng, D. and Rao, Y. (2014). Aggression and social experience: genetic analysis of visual circuit activity in the control of aggressiveness in Drosophila. Mol Brain 7, 55.

Rillich, J. and Stevenson, P. A. (2014). A fighter’s comeback: dopamine is necessary for recovery of aggression after social defeat in crickets. Hormones and behavior 66, 696–704.

Rillich, J. and Stevenson, P. A. (2015). Releasing stimuli and aggression in crickets: octopamine promotes escalation and maintenance but not initiation. Front Behav Neurosci 9.

Rollmann, S. M., Zwarts, L., Edwards, A. C., Yamamoto, A., Callaerts, P., Norga, K., Mackay, T. F. and Anholt, R. R. (2008). Pleiotropic effects of Drosophila neuralized on complex behaviors and brain structure. Genetics 179, 1327–36.

Rosay, P., Armstrong, J. D., Wang, Z. and Kaiser, K. (2001). Synchronized neural activity in the Drosophila memory centers and its modulation by amnesiac. Neuron 30, 759–70.

Siegel, R. W. and Hall, J. C. (1979). Conditioned responses in courtship behavior of normal and mutant Drosophila. Proc Natl Acad Sci USA 76, 3430–4.

Sitnik, N., Tokmacheva, E. and Savvateeva-Popova, E. (2003). The ability of Drosophila mutants with defects in the central complex and mushroom bodies to learn and form memories. Neurosci Behav Physiol 33, 67–71.

Stevenson, P. A., Dyakonova, V., Rillich, J. and Schildberger, K. (2005). Octopamine and experience-dependent modulation of aggression in crickets. J. Neurosci. 25, 1431–1441.

Tamura, T., Chiang, A. S., Ito, N., Liu, H. P., Horiuchi, J., Tully, T. and Saitoe, M. (2003). Aging specifically impairs amnesiac-dependent memory in Drosophila. Neuron 40, 1003–11.

Tan, Y., Yu, D., Pletting, J. and Davis, R. L. (2010). Gilgamesh Is Required for rutabaga-Independent Olfactory Learning in Drosophila. Neuron 67, 810–820.

Torroja, L., Chu, H., Kotovsky, I. and White, K. (1999). Neuronal overexpression of APPL, the Drosophila homologue of the amyloid precursor protein (APP), disrupts axonal transport. Curr Biol 9, 489–92.

Trannoy, S. and Kravitz, E. A. (2016). Strategy changes in subsequent fights as consequences of winning and losing in fruit fly fights. Fly, 0.

Trannoy, S., Penn, J., Lucey, K., Popovic, D. and Kravitz, E. A. (2016). Short and long-lasting behavioral consequences of agonistic encounters between male Drosophila melanogaster. Proc Natl Acad Sci USA 113, 4818–23.

Wang, L. and Anderson, D. J. (2010). Identification of an aggression-promoting pheromone and its receptor neurons in Drosophila. Nature 463, 227–31.

Wang, L., Dankert, H., Perona, P. and Anderson, D. (2008). A common genetic target for environmental and heritable influences on aggressiveness in Drosophila. Proc Natl Acad Sci USA 105, 5657–5663.

Wang, L., Han, X., Mehren, J., Hiroi, M., Billeter, J. C., Miyamoto, T., Amrein, H., Levine, J. D. and Anderson, D. J. (2011). Hierarchical chemosensory regulation of male-male social interactions in Drosophila. Nat Neurosci 14, 757–62.

Yoon, J., Matsuo, E., Yamada, D., Mizuno, H., Morimoto, T., Miyakawa, H., Kinoshita, S., Ishimoto, H. and Kamikouchi, A. (2013). Selectivity and plasticity in a sound-evoked male-male interaction in Drosophila. PLoS One 8, e74289.

Yu, D., Akalal, D. B. and Davis, R. L. (2006). Drosophila alpha/beta mushroom body neurons form a branch-specific, long-term cellular memory trace after spaced olfactory conditioning. Neuron 52, 845–55.

Yuan, Q., Song, Y., Yang, C. H., Jan, L. Y. and Jan, Y. N. (2014). Female contact modulates male aggression via a sexually dimorphic GABAergic circuit in Drosophila. Nat Neurosci 17, 81–8.

Yurkovic, A., Wang, O., Basu, A. C. and Kravitz, E. A. (2006). Learning and memory associated with aggression in Drosophila melanogaster. Proc Natl Acad Sci USA 103, 17519–24.

Zars, T., Fischer, M., Schulz, R. and Heisenberg, M. (2000a). Localization of a short-term memory in Drosophila. Science 288, 672–5.

Zars, T., Wolf, R., Davis, R. and Heisenberg, M. (2000b). Tissue-specific expression of a type I adenylyl cyclase rescues the rutabaga mutant memory defect: in search of the engram. Learn Mem 7, 18–31.

Zhou, C. and Rao, Y. (2008). A subset of octopaminergic neurons are important for Drosophila aggression. Nat Neurosci 11, 1059–67.

Zwarts, L., Versteven, M. and Callaerts, P. (2012). Genetics and neurobiology of aggression in Drosophila. Fly 6, 35–48.

